# Reconstructing time-resolved inter-residue distance distributions in a protein ensemble during functional dynamics in solution

**DOI:** 10.64898/2026.07.22.739050

**Authors:** Brad D. Price, Jackson Sheppard, Shiny Maity, Antonín Sojka, Joan-Emma Shea, Songi Han, Mark S. Sherwin

## Abstract

Reconstructing time-resolved inter-residue distance distributions during protein functional dynamics in the solution state is known to be a difficult and important problem. This article presents a technique for extracting spin-spin (as a proxy for residue-residue) distance distributions on doubly-spin-labeled proteins from rapid-scan time-resolved Gd-Gd electron paramagnetic resonance (rs-TiGGER) spectra recorded near room temperature in solution at 240 GHz. We use a best-fit technique that convolves a dipolar kernel matrix with an intrinsic, non-dipolar-broadened (single-labeled) spectrum. The kernel incorporates the effect of solution-state tumbling on the dipolar broadening using a correlation function that bridges the static and rapidly tumbling regimes. We apply the technique to *AsLOV2*, a protein domain with a dark-state crystal structure that is well-known from X-ray crystallography, but a less well-characterized and disordered tertiary structure that manifests after photoactivation at 450 nm. Informed by principal component analysis, we assume that the underlying distance distribution may be approximated by a sum of two Gaussian distributions. The fits returned time-resolved, light-activated populations with mean distances of 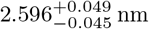 (dark) and 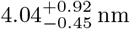 (lit) in the wild type, and 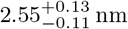 (dark) and 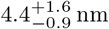 (lit) in an N414Q mutant, with nearly complete unfolding (within fit uncertainty) of the active, light-sensitive fraction. The extracted distance distributions and their accompanying uncertainties are consistent within uncertainty with molecular dynamics simulations of the equilibrated protein structure.

## 1 Introduction

Developing structural biology tools has been a focus of the biochemistry community for nearly a century [1]. Specifically, X-ray diffraction, nuclear magnetic resonance (NMR) and electron paramagnetic resonance (EPR) spectroscopy, and cryo-electron microscopy (cryo-EM) have all marked large leaps in the ability of scientists to determine the (static) structures of proteins. Very recently, the advent of AI-based protein structure prediction models has further accelerated this process, with the release of AlphaFold and other similar tools [2]. Mapping and understanding the structure of proteins allows for more precise determination of their function and therefore more precise design of drug targets [3, 4], as well as a better understanding of the underpinnings of all life on earth [5]. However, a well-known limitation of the majority of structure-function tools is that they require the protein to be in a non-native (frozen, crystallized, or otherwise immobilized) environment. Using the static structure of *AsLOV2* (PDB ID 2V1A, [6]) in Fig. 1 as an example—which was captured by X-ray crystallography—static structure techniques are particularly well-developed (more than 250,000 structures have been logged into PDB at the time of writing [7]), and are capable of providing structures with atomic resolution. However, there is a missing link to understanding protein function: the dynamic states of proteins in action remain unresolved with comparable precision. For example, published alongside the dark-state structure of *AsLOV2*, the lit state structure (PDB ID 2V1B) as recorded by X-ray crystallography, did *not* provide evidence for light-activated unfolding [8] even though it is known that unfolding takes place [9–11]: crystallization caused a restriction of the protein’s natural mechanical motion. To bridge this gap, progress toward time-resolved, solution-state tertiary structure measurements represents an important next frontier in structural biology.

**Fig. 1.**
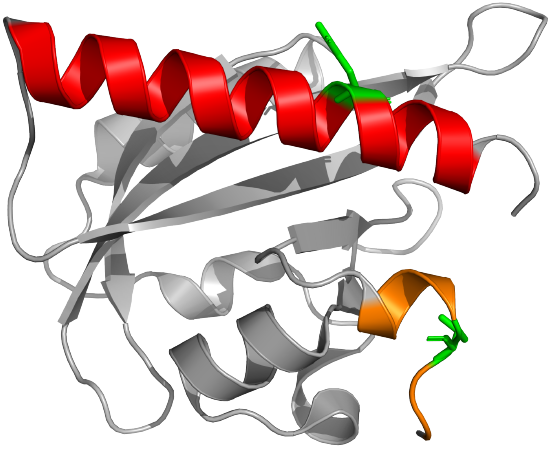
Dark-state structure of room temperature equilibrated *AsLOV2* (PDB ID 2V1A, [6, 8]). After undergoing blue light activation, it has been established that the J*α*-helix (red) becomes disordered, but it is unknown to what extent unfolding occurs or how the helix moves relative to the protein’s core. The protein’s A*α*-helix is shown in orange and the sites of the two spin labels, residue 537 on J*α* and residue 406 on A*α* are shown in green.

Tools for time-resolved protein structure measurements are undergoing rapid development. Time-resolved serial X-ray crystallography can yield internal protein kinematics with atomic spatial resolution and ps time resolution after a triggering event [12, 13], but cannot readily capture protein conformations with significantly disordered regions as these would not crystallize in their native form. Conformational ensembles captured by rapid freeze-quenching at variable delays after a triggering event can be studied by double electron-electron resonance (DEER) [14–16], solid-state NMR [17], and, more recently, cryo-electron microscopy (cryo-EM) [18]. In such methods, care must be taken to understand how the measured (frozen) conformational ensemble compares to the physiological-temperature ensemble of interest [18–20]. Solution state EPR [21–25], Förster Resonance Energy Transfer (FRET) [26–30], and time-resolved IR spectroscopy (*e*.*g*. [10, 31–33]) have made important contributions to time-resolved studies of protein structure, but also have limitations. Specifically, solution-state EPR, at least to date, has been primarily limited to studies of local (secondary) structure through correlating the resonance linewidth to the relative mobility of the label in specific local conformational states (*e*.*g*., the pioneering work by Steinhoff and Hubbell on bacteriorhodopsin unfolding, [21]). Time-resolved IR spectroscopy has sub-ps time resolution, but faces a similar drawback: it is limited to studying the local, secondary structure dynamics of specific chemical bonds [10, 34]. FRET, on the other hand, especially single-molecule FRET, is remarkably precise and informative on time-resolved tertiary structure. However, it is often limited to a mean distance ⟨*r*⟩ because the fluorescence carries no spectral information, and thus does not inform on distance *distributions* of an ensemble, making it most powerful for measurements of single molecules or average conformations. There are significant (ongoing) efforts to extract inter-residue distance distributions via extensive cross-validation with *a priori* knowledge of *P* (*r*), including refs. [28, 35–38].

We previously introduced Time-resolved Gd-Gd EPR (TiGGER) to resolve transient spin-spin (residue-residue) dynamics in a solution-state protein ensemble [39] on a model protein, *AsLOV2* : a light-activated phototropin 1 domain that has a well-known *Jα*-helix undocking and subsequent unfolding after light activation [9, 40] (absorption peaks in the blue (450 nm) range [41, 42]). *AsLOV2* remains of interest and details of its photocycle are still being uncovered [43–46]. Additionally, *AsLOV2* has applications in optogenetics, providing a pathway to nanometer-scale photoswitches (*e*.*g*. [47, 48]), an application for which solution-state, time-resolved structure measurements may be particularly important. However, at the time of our previous reporting, we had yet to recover spin-spin distances from TiGGER data. Because TiGGER (based on rapid-scan EPR) records the entire EPR lineshape of interest as a function of time, the full set of relevant spin-spin couplings is readily available, and the task of recovering distance distributions from the lineshape is only a numerical one. In this article, we present a technique that uses Pake-convolution-based forward simulation to artificially broaden the second moment of a singly-labeled EPR spectrum and generate a best fit to the experimentally recorded doubly-labeled spectrum. Complementary to other structural biology probes, with the addition of the numerical method described in this paper, TiGGER provides a method to reconstruct the distribution of spin-spin distances of a time-resolved conformational change in real-time and the solution state.

As a brief recap to the working principle of TiGGER, spin-spin dipolar coupling results from the magnetic moment of a spin *Ŝ*_2_ imparting a small perturbation to an applied static field on another spin, *Ŝ*_1_. The contribution of these effects to the electron spin Hamiltonian is the following:

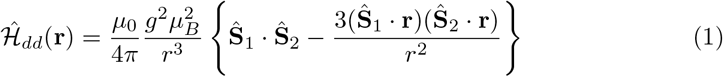

where *µ*_0_ is the permeability of free space, *g* is the free electron *g*-factor (*g* ≈ 2), *µ*_*B*_ is the Bohr magneton, ***Ŝ***_*i*_ represents a spin operator for an electron *i*, and **r** is the vector spanning the space between the two spins. Following the simplification from Ch. IV of Abragam [49] and collecting the relevant “secular” terms, 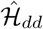 can be written

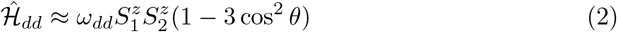

where *θ* is the angle between the two spins and the applied magnetic field, and *ω*_*dd*_ is the dipolar coupling strength as a function of spin-spin distance, *r*:

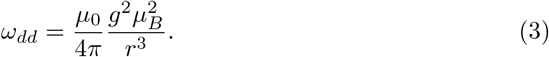

For unlike (distinguishable) spins, the resonance frequencies go from (*ω*_1_, *ω*_2_) to (*ω*_1_ ± *ω*_*dd*_*/*2) in the weak coupling regime (assuming *ω*_1_ ≈ *ω*_2_ ≪ *ω*_Zeeman_) which results in a line splitting (or broadening, in case *ω*_*dd*_ ≪ Δ*ω*_intrinsic_) by an amount *ω*_*dd*_. The problem to solve, then, is recovering the magnitude of spin-spin dipolar coupling and thus the underlying distance (distribution) of the ensemble from the broadened dataset. This article presents a technique for backing out these distributions via calculating global best-fit parameters from a forward-simulated Pake convolution. The results of a representative time-resolved experiment on *AsLOV2* are presented in Section 3.

## 2 Pake convolutional fitting

To perform the fit, a Pake convolution technique was developed that relies on adding the effect of dipolar spin-spin coupling to an otherwise identical spectrum that is not dipolar broadened [50–52], which can be achieved by only spin labeling one residue. By this method, the effect of the dipolar coupling may be represented entirely as a convolution between the unbroadened spectrum as a function of field (*B*), *S*(*B*) with two-spin dipolar coupling *D*(*B*):

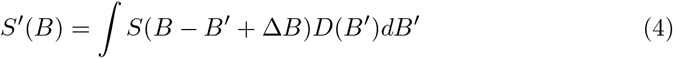

where *S*^*′*^(*B*) is the observed, broadened lineshape, and Δ*B* is a field shift that can correct small offsets between resonance positions of broadened and unbroadened datasets. When immobilized, the coupled spectrum is typically called a “Pake pattern”. This term is not used here, as the solution-state broadening is not explicitly a Pake pattern, though it is analogous. To simplify the generation of the dipolar patterns that govern the line, it is important to reduce the space of acceptable distance distributions to make the fit robust. To inform this reduction, we performed principal component analysis (PCA) to determine the number of changing spectral components during a light-activated TiGGER experiment.

The results of the PCA analysis are shown in Fig. 2; here, *AsLOV2* was spin labeled at sites 406 and 537 (shown green in Fig. 1). The sites 406 and 537 were chosen because 406 is on the N-terminus, near the core of the protein, and 537 is on the C-terminus, on the J*α*-helix that is known to become disordered upon light activation [10]. The C-terminus was expected to move away from the protein after unfolding, as the well-defined structure that constrains it near the core was eliminated with light activation, making it an ideal model system to demonstrate convolutional distance distribution reconstruction.

**Fig. 2.**
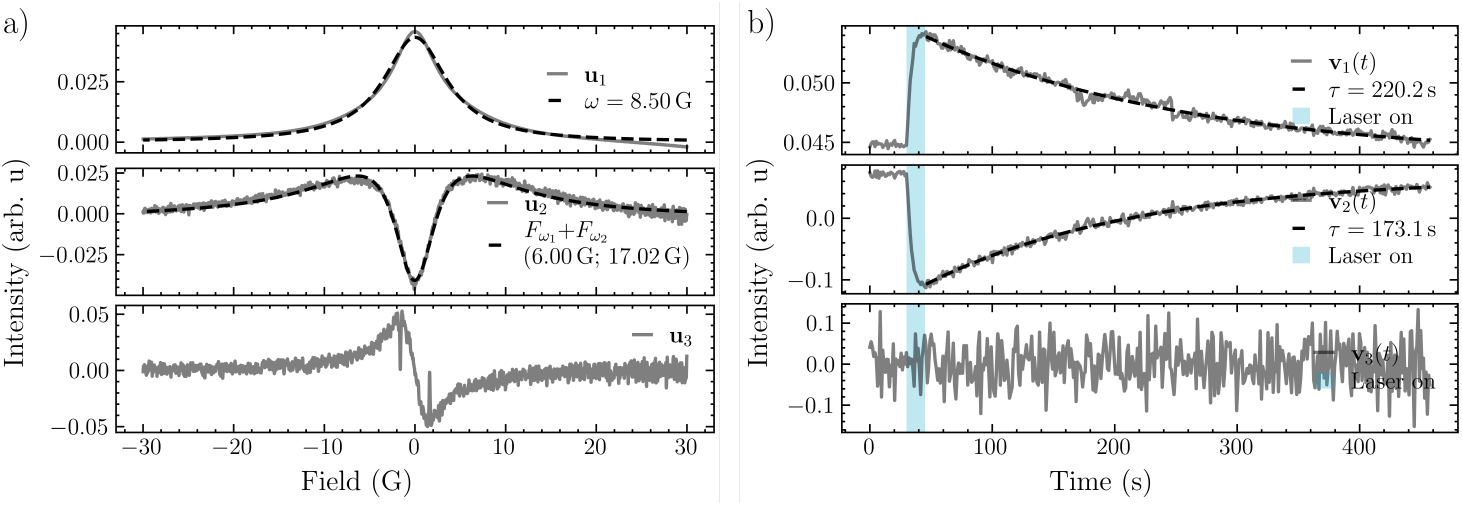
PCA results for the 2D time-and-field resolved TiGGER data of WT *AsLOV2* spin-labeled at sites 406 and 537 and at 10 ^°^C. Temperature was lower than 20 ^°^C for this experiment because it improved SNR and therefore made the principal components easier to interpret. (a) The principal components of the spectra. The first and second components are fitted with a Lorentzian lineshape and the difference of two Lorentzian lineshapes, respectively. (b) The temporal weight of each PCA component. The first and second coefficients are fitted to decaying exponentials that are related to the re-folding (and re-broadening) dynamics of *AsLOV2*. Vertical blue bar marks the times that blue light illumination was on.

As seen in the middle panels of Fig. 2a and 2b, there appear to be two line-shape components that flip-flop when the laser was turned on. Specifically, the time-dependence of the second component was initially positive and flips negative. PCA was performed on the raw data (not mean-centered), because it made the results easier to interpret; the first component was related to the mean spectrum, leaving the majority of the variance to the second component. As can be seen in Fig. 2, the second component itself was made up of a broad (positive) component and a narrow (negative) one. When the coefficient flipped during laser illumination (shown by the vertical blue bar), the broad contribution shrunk and the narrow contribution grew. This implied that the accessible distance distribution may be sufficiently modeled by a sum of two Gaussians, with one near (strongly dipolar broadened) and one far (weakly dipolar broadened) component.

Informed by the PCA, we chose to represent the distance distribution *P* (*r*) as a normalized sum of two Gaussian distributions, with fixed widths and means, and exponentially time-varying amplitudes:

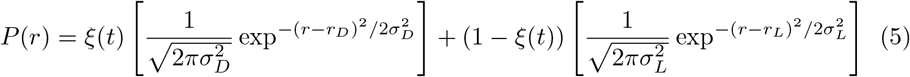

where

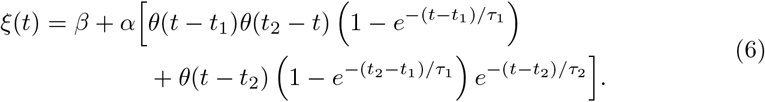

Here, *β* represents the fraction of protein that is unfolded in the dark state (and has no time dependence) and *α* represents the fraction of proteins that respond to light activation. *ξ*(*t*) must therefore be bounded between zero and one, and since the term in parentheses in Eqn. 6 is less than or equal to one, the sum of *α* and *β* must also be less than or equal to 1. Note that subscript in equation (5), *D* is for dark (folded), and *L* is for lit (unfolded). In equation (6), *t*_1_ is when the blue light is turned on, *t*_2_ is when it is turned off. *τ*_1_ and *τ*_2_ are the exponential lifetimes of unfolding and refolding, respectively. Gaussians were chosen as the sub-population distribution as a result of the central limit theorem—a large number of randomly-sampled, equally likely orientations for both the protein and spin label ligands tend toward normal distributions in large numbers (1 *µ*L ∗ 1 mM ∼ 10^15^ here) [53].

An implicit assumption of the model represented by equation (5) is that the individual proteins within the ensemble may only be in one of two states: folded or unfolded. This implies that there are few-to-no long-lived (stable or semi-stable) intermediate states of the photocycle, and that the transition proceeds rapidly with respect to the timescale of the experiment. On an individual protein level, it is expected that this is the case (individual unfolds/folds expected to be ∼ *µ*s; see, *e*.*g*. [54]). Without this assumption, each intermediate state would require its own distance distribution and lifetime, greatly complicating the fit and its interpretation.

The dipolar kernel, *D*(*B*) in equation (4), is analogous to a Pake pattern. Because the sample was in solution at room temperature, a typical dipolar Pake (“powder”) pattern is unreasonably sharp given the Brownian motion of the ensemble and other line broadening relaxation mechanisms [55]. The kernel, then, may be written as the Fourier transform of Brownian diffusion correlation function *G*(*t*) that preserves the second moment of the full Pake pattern [49, 56]

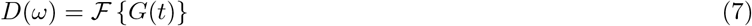

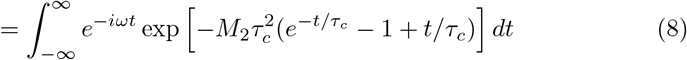

A change of variables from frequency to field units, *ω* = *γB*, gives the field-domain kernel *D*(*B*) used in Eqn. 4, where *M*_2_ is the second moment of the spin-spin dipolar splitting. It can be seen that for 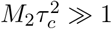, the lineshape collapses to a Gaussian with width 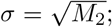 if *M*_2_*τ*_*c*_ ≪ 1, it collapses to a Lorentzian with width Γ = *M*_2_*τ*_*c*_. To calculate *M*_2_, we impose a few assumptions:

1. Gd-sTPATCN, the spin label used in this work, is known to behave approximately as a spin-1/2 due to large ZFS coupling smearing non-central transitions into a broad baseline that is difficult to resolve with EPR [57, 58].
2. The “sensor” spin, therefore, can be treated as a spin-1/2 in the neighborhood of a spin-7/2. Since the *S* = 7*/*2 spins have a broad distribution of resonance fields due to ZFS strain and the direction of individual ZFS tensors, the weak coupling regime is a valid approximation for almost all spin pairs.
3. At room temperature and 8.6 T, the *S* = 7*/*2 manifold is approximately evenly populated.
4. The coupling operator of interest is *Ŝ*_*z*_ , as that is the term of the magnetic moment that splits the 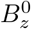 resonance.
5. The angle coupling term may be spatially averaged over 4*π*, valid for globular proteins tumbling in solution and consistent with Abragam’s treatment of nuclear spin interactions in isotropic motion [49], which states that “The second moment of the absorption curve is unaffected by narrowing ‘motion’.”

Therefore,

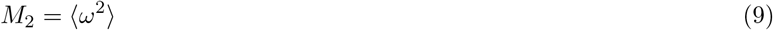

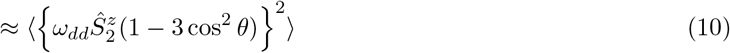

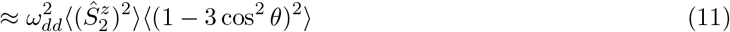

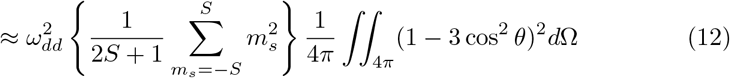

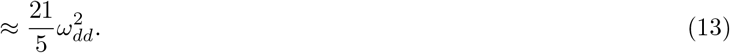

Crucially, for spin-spin distance extraction to work, dipolar coupling must survive the effect of motional narrowing. As shown in a previous publication [39], *some* dipolar coupling does survive. For two spin-1/2 electrons, *ω*_*dd*_(2.8 nm) ≈ 2*π* · 2.3 MHz; an estimate for the protein’s rotational correlation time, *τ*_*c*_, based on the radius of gyration, *R*_*g*_, of crystal structure *AsLOV2* (PDB 2V1A [6, 8]), is approximately *τ*_*c*_ = 13 ns, assuming the radius of hydration, *R*_*h*_ = 0.775*R*_*g*_, for a compact, folded protein, and Stokes-Einstein hard sphere rotational diffusion [59, 60]. *R*_*g*_ was calculated by appending the spin labels to the protein PDB using `chilife` and then `MDAnalysis`’s `radius_of_gyration()` function [61–64]. A correlation time in this range results in *ω*_*dd*_*τ*_*c*_ ≈ 0.2, squarely in the intermediate motion regime, justifying the use of the correlation function in Eqn. 8 that bridges the two regimes.

As seen in Clayton *et al*. [65], Gd(III) spin labels are capable of resolving *r*^−3^ dipolar broadening in distances above 3 nm. However, that article only presents results from either frozen or otherwise immobilized samples, restricting motional narrowing, which limits the application to solution-state results. Here, we make the assumption that their experiment using room-temperature immobilization (in a glassy trehalose matrix) only differs in rotational correlation time from what is presented here. Therefore, because convolution simply sums second moments, 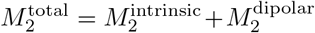 any intrinsic (isotropic, not affected by rotation) broadening effects are included in 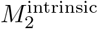 via the convolution, and the effect of tumbling on the dipolar component of the lineshape is included in 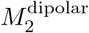, resulting in a complete treatment of the lineshape. We also note here that Eqn. 8 explicitly treats the crossover from *r*^−3^ broadening to *r*^−6^ broadening, as the linewidth in the Gaussian (static, inhomogeneously broadened) regime scales as 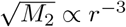 and in the Lorentzian (relaxation-broadening, homogeneous) one as *M*_2_ ∝ *r*^−6^ [66].

## 3 Results

To demonstrate this technique, experiments were performed on wild type (WT) and N414Q-mutated *AsLOV2*. The N414Q mutation was chosen because it is known to slow the timescale of light-induced motion, but the higher-order effects that cause the slowing are actively being investigated [46].

To benchmark the fits, molecular dynamics (MD) simulations were performed. It is challenging, at this time, for MD to simulate directly the conformational changes that result from quantum mechanical processes involved in photoexciting the FMN chromophore in AsLOV2. However, as demonstrated in [46], pressure does a remarkable job of replicating the effects of light activation in *AsLOV2*, as evidenced by DEER and MD simulations. Thus MD simulations were performed at 1 bar for the equilibrium dark state dynamics, and 3 kbar as a proxy for the lit state dynamics. Specifically, the simulations began with the crystal structure of dark-state *AsLOV2* (PDB ID: 2V1A) that was allowed to equilibrate to ambient and high-pressure conditions using the implicit solvent protocol outlined in [67]. Representative conformations were then equilibrated in explicit solvent and subjected to 500 ns of conventional MD. Then, the resulting set of conformations from the 200-500 ns window were used as independent protein structures for `chilife`, a software package for calculating spin-spin distances of site-directed spin labeled PDB structures [62]. From these, the distribution of distances for each environment was calculated, including the full space of conformations accessible to both the proteins and the attached Gd-sTPATCN spin labels, using the off-rotamer sampling tool provided by `chilife` [61](*N*_samples_ = 5000). There are additional details about the MD methods in S.I. Sec. S1 and [46]. Unfolding of the dark state structure at high pressure (3 kbar) is treated as a proxy for the light-induced unfolding studied by TiGGER. The results of the calculations are presented in Figs. 3 and 4.

**Fig. 3.**
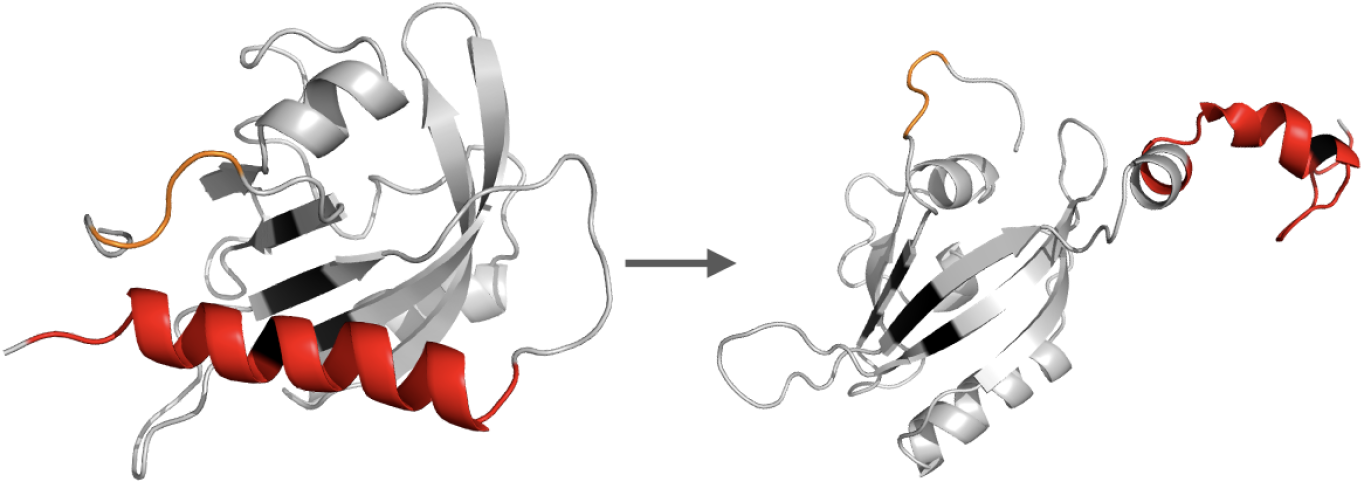
High-pressure (3 kbar) equilibrated MD conformations demonstrating pressure-induced unfolding of the J*α*-helix (red), simulating what is expected to occur under light activation. Left structure shows *t* = 0 and right structure shows simulation time *t* = 200 ns. Simulation was conducted in AMBER with the PLUMED SASA plugin.

**Fig. 4.**
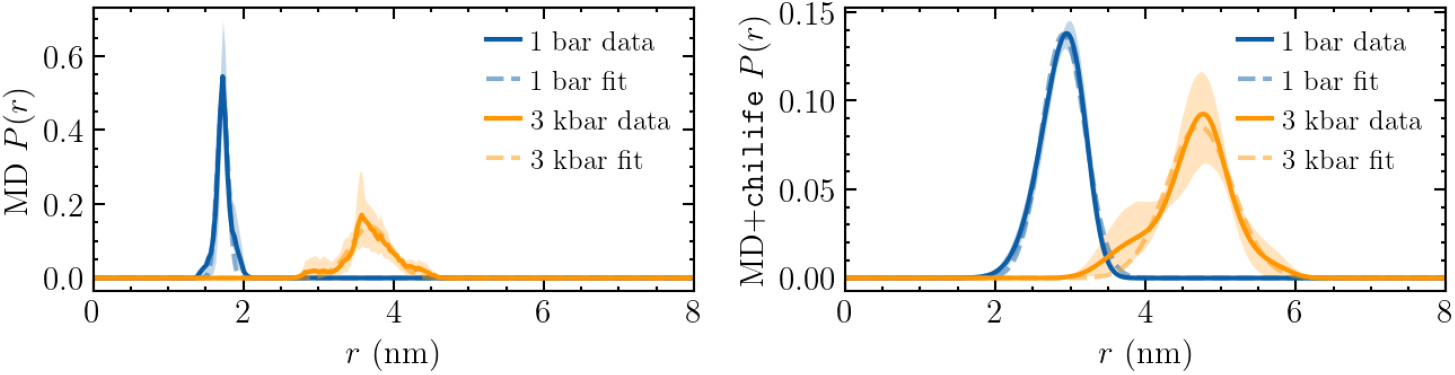
Results of calculated residue 406-537 distances resulting from MD simulations that sampled the whole 300 ns production trajectory without (left) and with (right) attached spin labels. 1 bar, ambient, dark-state equilibrated structures are shown in blue and 3 kbar simulated lit-state structures are shown in orange along with their accompanying Gaussian fits (shown with dashed lines, fitted using `scipy.optimize.curve fit()`). 95% confidence intervals for the distributions, calculated with block bootstrapping (block size of 49 ns, the largest autocorrelation time across the four datasets) are shown with the transparent bands. MD analysis (no spin labels) returned *µ* = 1.725 ± 0.001 nm, *σ* = 0.073 ± 0.001 nm (raw mean ⟨*r*⟩ = 1.732 nm) at 1 bar and *µ* = 3.688 ± 0.005 nm, *σ* = 0.264 ± 0.005 nm (raw mean ⟨*r*⟩ = 3.695 nm) at 3 kbar. MD+`chilife` best fits returned *µ* = 2.907 ± 0.002 nm, *σ* = 0.289 ± 0.002 nm (raw mean 2.875 nm) at 1 bar and *µ* = 4.714 ± 0.006 nm, *σ* = 0.439 ± 0.006 nm (raw mean 4.640 nm) at 3 kbar. The volume accessible to the Gd(III) spin labels increase the spin-spin distances by roughly 1 nm while smoothing and broadening the distributions.

The Pake convolutional fits were performed using the `lmfit` python library [68] using a basinhopping technique, designed to broadly sample the parameter space to find a global minimum, and then further optimized locally with a single run of a tighter tolerance least-squares routine. Because the model has ten fitted parameters and the solution is not unique, it is necessary to constrain the fit parameters to obtain a physically reasonable result. In order to achieve this, a select few parameters were provided log-normal Gaussian priors that nudged the fit in the direction of results that were estimated *a priori*. These priors were of the form

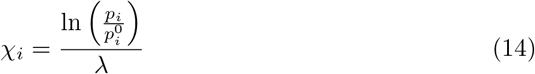

where *χ*_*i*_ was squared and added to the total *χ*^2^. To provide a scale for the priors, *χ*^2^ was normalized by the number of fit points (*N* ^field^ ∗ *N* ^time^) and the standard deviation of the noise, which was estimated by subtracting a smoothed (Gaussian filtered) version of each time slice from itself. The width of the Gaussian filter *σ* was calculated using an *L*-curve to find the value that produced the greatest Menger curvature [69–71] in a plot of noise vs. *σ* values (see S.I. Fig. S1). This calculation assumes that the signal and noise can be separated according to their frequencies, with signal at low frequencies and noise at high frequencies. This approximation is validated by the large number of spins in the ensemble with distributed local fields effectively smoothing any high-frequency structure in the signal; further, noise near DC should be effectively eliminated by the coherent detection and baseline subtraction scheme outlined in [39]. As seen in S.I. Fig. S2, there was a portion of high-frequency signal at the peak of the EPR spectra. In order to only select the region with slowly-varying lineshapes and thus return a signal-smoothed result that was purely noise, the data was masked for |*B*| *>* 10 G .

An *L*-curve was also generated to find the crossover point of maximum Menger curvature in order to balance the contributions between minimizing the *χ*^2^ of the fit to the data and agreeing with the set of priors (see S.I. Fig. S3). For the WT and N414Q samples, the optimal *λ* was calculated to be *λ* = 0.927 and *λ* = 0.534, respectively. The fit parameter constraints and priors values are presented in Table 1:

**Table 1.** Parameters, constraints, and priors for Pake convolution-based TiGGER fits. Total experiment duration represented by *t*_max_.

| Parameter | Units | Symbol | Fit Range | Prior |
| --- | --- | --- | --- | --- |
| Scale coefficient | — | $A$ | [0.75, 1.25] | — |
| Unfolding time | s | $\tau_1$ | $[0.002t_{\text{max}}, 0.02t_{\text{max}}]$ | — |
| Refolding time | s | $\tau_2$ | $[0.05t_{\text{max}}, 0.5t_{\text{max}}]$ | Linewidth $\tau$ [39] |
| Unfolded dark-state fraction | — | $\beta$ | [0.0, 0.4] | 0.06 [72] |
| Active fraction | — | $\alpha$ | $[0, 1 - \beta]$ | — |
| Dark-state $\langle r \rangle$ | nm | $r_D$ | [2.0, 4.5] | — |
| Dark-state variance | nm | $\sigma_D$ | [0.1, 2.0] | 0.29 (Fig. 4) |
| Lit-state $\langle r \rangle$ | nm | $r_L$ | $[r_D + 0.5, r_D + 3.0]$ | — |
| Lit-state variance | nm | $\sigma_L$ | [0.2, 2.0] | 0.44 (Fig. 4) |
| Field axis shift | G | $\Delta B$ | [-0.25, 0.25] | — |

To minimize the effect of these priors on the result of interest—primarily, the spin-spin distances throughout the protein unfolding and refolding—priors were only provided for variables that we had independent estimates for and that did not directly inform the output distances. These parameters were (1) the re-folding time, *τ*_2_, as that value should be close to the relaxation time of the width of the EPR line as reported in [39]; (2) the fraction of proteins that are unfolded in the dark, informed by [72] at 6%; and (3) the width of the two spin-spin distance distributions, *σ*_1_ and *σ*_2_, informed by the MD-and-`chilife` simulations of equilibrated *AsLOV2* shown in Fig. (4).

The fit residue was windowed to select regions of the dataset that highlighted the kinetics rather than over-weighting the fit with pure noise (in the wings of the field domain) or primarily static data, at long times after the laser was turned off at *t*_2_. Specifically, the field domain was windowed with a super-Gaussian in order to highlight the center of the field axis while reducing the contribution to the fit *χ*^2^ from the wings using

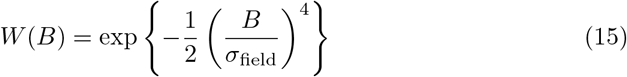

with *σ*_field_ = 15 G (see S.I. Fig. S4). The time-axis data was windowed with an exponential decay that peaks at *t*_2_, when the laser was turned off:

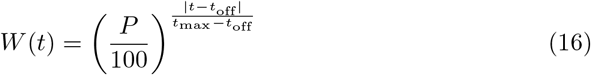

here, *P* is the desired weight given to the time-axis at the maximum time point. For this article, *P* = 10 (window shown in S.I. Fig. S5).

Results of best-fits for WT and N414Q time-and-field data matrices are shown in Fig. 5 and example temporal slice at *t* = *t*_2_ is shown in Fig. 6. These fits returned the following values for the parameters presented in Table 1 above.

**Fig. 5.**
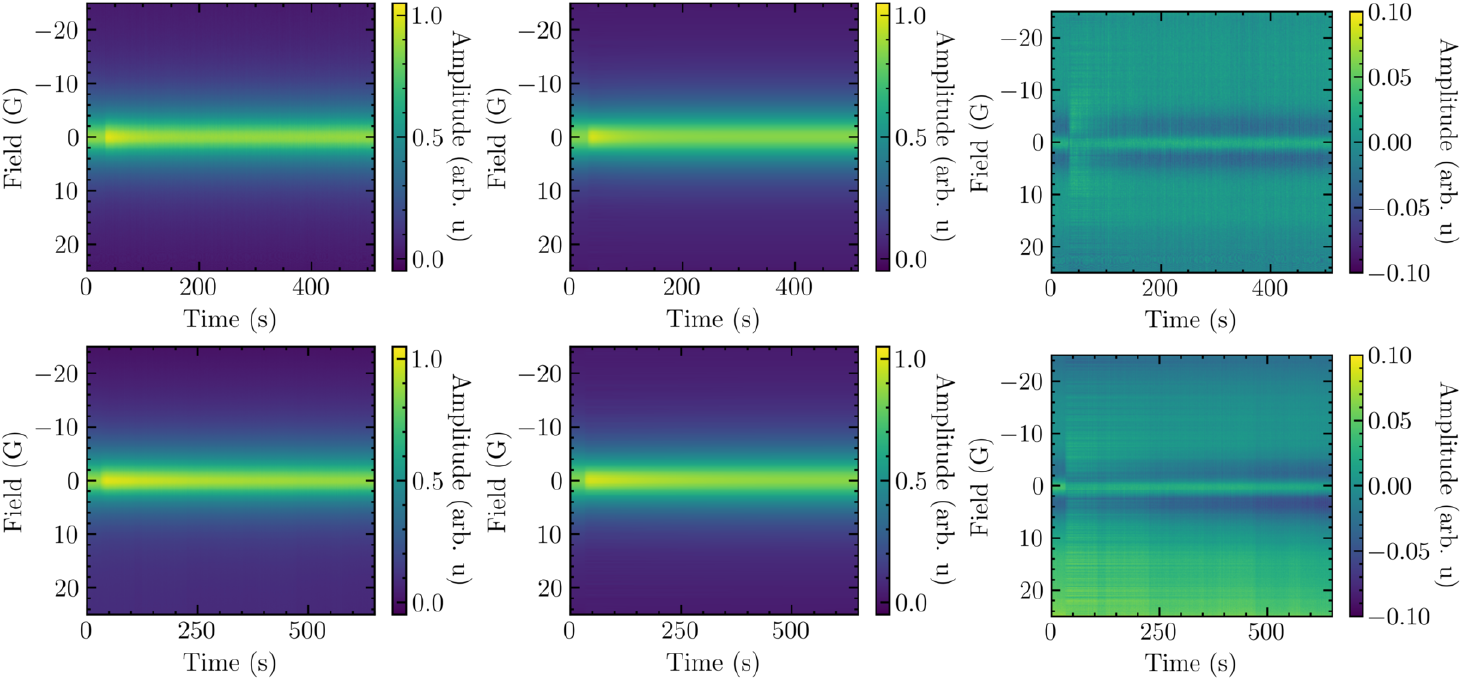
Comparison of field-and-time resolved data (left column) to Pake convolution-generated fits (middle column) for both wild-type (top) and N414Q mutant (bottom) *AsLOV2*. Residues for each fit (right) show roughly 5% or less deviation between data and fit. For the N414Q experiments, the laser was left on for additional time to ensure maximum narrowing: *t*_2_ − *t*_1_ = 15 s rather than the 10 s for the WT. Both experiments were performed at ∼ 20 ^°^C.

**Fig. 6.**
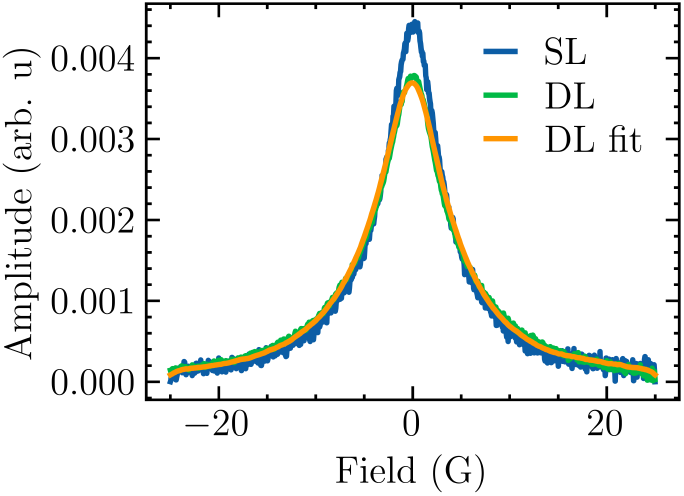
Temporal slice at *t* = *t*_2_ for wild-type single label (unbroadened, blue), double label (green), and the convolved best fit (orange).

The 95% confidence intervals were calculated using a profile likelihood method: each parameter was fixed at a series of values spanning the acceptable range, all remaining free parameters were re-optimized, and the minimum *χ*^2^ at each fixed value was recorded. The width of the resulting *χ*^2^ profile determines the CI (example profiles shown in S.I. Fig. S7; discussed in detail in S.I. S3).

**Fig. 7.**
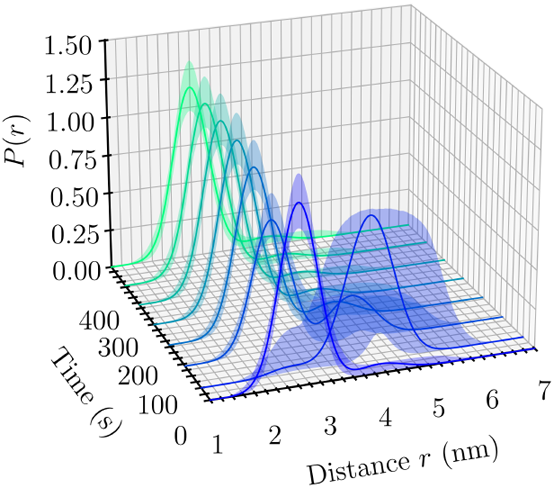
Summed Gaussian distance distributions as returned by the best fit of the model to the TiGGER data of WT *AsLOV2* at 18 ^°^C. Eight of roughly 500 frames are shown. The sample was illuminated between frames one and two. Light-activated unfolding and refolding is shown, resulting in a large shift in relative near/far populations. See S.I. Fig. S8 for equivalent N414Q figure. Confidence intervals were generated by Monte Carlo sampling skew-asymmetric Gaussian distributions with the 95% CI as 1.96*σ* [73] (see S.I. Fig. S9).

## 4 Discussion and conclusion

The results of the TiGGER distance determination method on *AsLOV2* are consistent within uncertainty with the results of the MD simulations and with the well-known effect of J*α*-helix unfolding upon blue light activation (Table 2) [25, 32, 39, 46, 72]. The Pake convolutional best-fit forward simulation technique is able to provide reasonable values for the summed Gaussian distance distribution model, highlighting a fairly rapid (*<* 5 s) unfolding and slower (∼ 55 s) refolding in the wild type, and slower (∼ 5 s) unfolding and (∼ 220 s) refolding in the N414Q mutant. The 95% confidence intervals for the fitted parameters are quite narrow, particularly for parameters that strongly inform the fit quality (*e*.*g. A, α, β, r*_*D*_); the CIs are wider for parameters that more weakly determine the fit (*e*.*g. τ*_1_, *r*_*L*_). This was expected, as *τ*_1_ makes up a very small fraction of total data in the fitted matrix (15 s*/*500 s ≈ 3%), and because electron-electron dipolar broadening falls off extremely quickly with distance, making distances above ∼ 4 nm nearly impossible to distinguish (see [65], for example). It is worth noting that the results here—though *β* was informed by prior publications—were in agreement with near-total unfolding under blue light illumination that has been reported by others at 5.98 ± 0.08% dark-state-unfolded (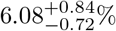 here) and 91 ± 6% lit-state-unfolded (99_−16_% here, where † denotes a CI bound not resolved within the parameter’s acceptable range) [72]. The light-responsive fraction, 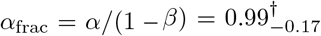 (confidence interval for *α*_frac_ was profiled independently, see S.I. Fig. S7), is consistent with near-complete photo-response among light-sensitive (initially folded) proteins; the upper CI bound is not resolved as *α*_frac_ ≤ 1 is a hard physical constraint. There is also some evidence resulting from the fits that the N414Q mutation does indeed increase the extent of motion 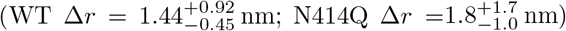, though the error bars on *r*_*L*_ are very large—additional studies on this effect are required before conclusions may be made. Additionally, N414Q may reduce the fraction of light-active proteins, as 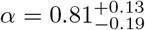, which is more than 10% less than the WT case; however, the CI for this parameter represents a very large uncertainty, including even the maximum *α* = 1 − *β* = 0.94.

**Table 2.** Best-fit values and 95% confidence intervals (CIs) for physical parameters, calculated via profile likelihood (see S.I. Fig. S7). Superscript and subscript offsets give the upper and lower CI bounds relative to the best-fit value. The MD column gives Gaussian parameters fitted to the dark (1 bar) and lit (3 kbar, proxy for illuminated) state distance distributions from MD simulation of *AsLOV2* (Fig. 4); *σ* priors were informed by these fitted MD values.

| Parameter | WT | N414Q | MD | Units |
| --- | --- | --- | --- | --- |
| $A$ | $1.0513^{+0.0065}_{-0.0060}$ | $1.067^{+0.015}_{-0.013}$ | — | — |
| $\tau_1$ | $2.7^{+2.3}_{-1.7}$ | $4.6^{+8.4}_{-4.5}$ | — | s |
| $\tau_2$ | $55.4^{+7.2}_{-6.2}$ | $221^{+66}_{-50}$ | — | s |
| $\beta$ | $0.0608^{+0.0084}_{-0.0072}$ | $0.061^{+0.020}_{-0.015}$ | — | — |
| $\alpha$ | $0.93^{+0.01}_{-0.16}$ | $0.81^{+0.13}_{-0.19}$ | — | — |
| $r_D$ | $2.596^{+0.049}_{-0.045}$ | $2.55^{+0.13}_{-0.11}$ | $2.9074 \pm 0.0037$ | nm |
| $\sigma_D$ | $0.348^{+0.054}_{-0.048}$ | $0.40^{+0.15}_{-0.12}$ | $0.2888 \pm 0.0037$ | nm |
| $r_L$ | $4.04^{+0.92}_{-0.45}$ | $4.4^{+1.6}_{-0.9}$ | $4.714 \pm 0.011$ | nm |
| $\sigma_L$ | $0.439^{+0.060}_{-0.052}$ | $0.44^{+0.14}_{-0.11}$ | $0.439 \pm 0.011$ | nm |
| $\Delta B$ | $0.0277^{+0.0046}_{-0.0045}$ | $0.0323^{+0.0088}_{-0.0086}$ | — | G |

This article presents progress toward a technique that is capable of extracting time-resolved distance distributions from an ensemble of solution-state, room temperature protein domains undergoing functional dynamics. The technique makes use of the uniquely narrow Gd-sTPATCN lineshape and sensitivity to spin-spin broadening at high field to extract spin-spin (as a proxy for residue-residue) distance distributions. The dipolar broadening model is remarkably simple, treating only the effect of room temperature motion on spatially averaged spin-spin coupling, and is still able to return results consistent with MD simulations while only requiring a few priors to stabilize the fit results. Though similar measurements may be made with other techniques, to the knowledge of the authors, none are capable of directly reconstructing distance distributions under life-like conditions, making TiGGER a useful complement to a range of other well-established techniques. As presented here, a combination of structural biology tools (in this case, high-field EPR, X-ray diffraction, and molecular dynamics) can be used to drive an iterative process that converges on a more precise result than any single technique can provide on its own.

## Supplementary information

A PDF document containing the supplementary information and figures referenced in this article is available online.

## Declarations

### 4.1 Funding

This work was supported by the National Science Foundation under MCB-2025860. Use was made of computational facilities purchased with funds from the National Science Foundation (CNS-1725797) and administered by the Center for Scientific Computing (CSC). The CSC is supported by the California NanoSystems Institute and the Materials Research Science and Engineering Center (MRSEC; NSF DMR 2308708) at UC Santa Barbara.

### 4.2 Declaration of competing interests

The authors declare no competing interests.

### 4.3 Data and code availability

TiGGER distance-distribution-reconstruction data and code are available at https://doi.org/10.5281/zenodo.21251338. `chitraj`, MD+`chilife` code is available at https://doi.org/10.5281/zenodo.21422884.

### 4.4 Declaration of generative AI and AI-assisted technologies in the manuscript preparation process

During the preparation of this work, the authors used Anthropic’s Claude Code model Sonnet 4.6 and 5. It was used to iterate on, comment, and improve software for speed and reliability. It was also used to scan the manuscript for spelling, clarity, and equation-formatting errors. After using this tool, the authors reviewed and edited the content as needed and take full responsibility for the content of the published article.

## Notes

https://doi.org/10.5281/zenodo.21251338

https://doi.org/10.5281/zenodo.21422884

### Competing Interest Statement

The authors have declared no competing interest.

