## Supplementary Information pdf for "Reconstructing time-resolved inter-residue distance distributions in a protein ensemble during functional dynamics in solution"

### Supplementary Information: Reconstructing time-resolved distance distributions after activation of a protein ensemble at room temperature in solution

Brad D. Price, Jackson Sheppard, Shiny Maity, Antonín Sojka, Joan-Emma Shea, Songi Han, Mark S. Sherwin

#### S1 Molecular dynamics details

Lit-state conformations of *AsLOV2* for `chilife` analysis were generated using a two-phase MD protocol.

In the first phase, we followed the protocol of [1] to sample at 3 kbar. This system and an ambient-pressure (1 bar) control were each simulated for 200 ns at 298 K in AMBER [2] using the PLUMED SASA implicit-solvent plugin [1, 3]. For each pressure condition, conformations were clustered by RMSD using the GROMOS [4] algorithm in GROMACS [5] with a 0.4 nm cutoff. The centroid of the most populated cluster was carried forward.

In the second phase, these centroids were prepared in GROMACS [5] using the AMBER99SB-ILDN force field [6] with the TIP4P-Ew water model [7]. Each centroid was placed in a dodecahedral simulation box with a minimum solute-edge distance of 1.0 nm, solvated, and neutralized with four $Na^+$ ions. Systems were energy-minimized until the maximum force fell below 1000 kJ mol$^{-1}$ nm$^{-1}$. All simulations used the leap-frog integrator [8] with a 2 fs time step. Bonds involving hydrogen were constrained with LINCS [9] and water geometries with SETTLE [10]. Lennard-Jones interactions were truncated at 1.0 nm, and electrostatics were treated with particle-mesh Ewald (PME) [11] using a 1.0 nm real-space cutoff, 0.16 nm grid spacing, and fourth-order interpolation. Equilibration comprised 5 ns with the Berendsen thermostat and barostat [12] followed by 5 ns with the velocity-rescale thermostat [13] and Parrinello-Rahman barostat [14], both at 300 K and at the pressure of the corresponding first-phase condition (1 bar or 3 kbar). Production runs were 500 ns with coordinates saved every 10 ps, and analysis was restricted to the 200-500 ns window. Frames from these trajectories served as independent protein structures input to `chilife` off-rotamer sampling with 5000 samples [15], as described in the main text.

### S2 Fitting model details

To minimize human intervention to the fit processes, $L$-curves were used to optimize the value of the smoothing width, $\sigma$, and the strength of the fit priors, $\lambda$. The region of maximum Menger curvature [16–18] of the $L$-curve was used for both, as it selects the region where contributions from the two regimes both have an effect on the resulting metric (*e.g.* noise $\sigma$ in Fig. S1 or fit quality $\chi^2$ in Fig. S3).

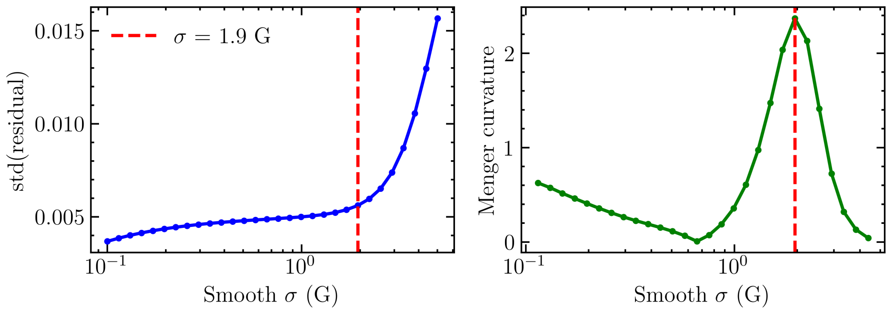


**Fig. S1** (Left) Noise $\sigma$ $L$-curve highlighting the Gaussian noise figure required to optimally smooth the data. Smaller $\sigma$s tend toward no noise, and larger $\sigma$s under-fit the EPR spectrum shape leaving large residuals. (Right) Menger curvature highlighting the point of maximum curvature between the two regimes; found to be 1.9 G for the WT dataset.

Noise was only estimated in the wings of the field profiles, as high-frequency signal regularly occurred near the peak of the EPR spectrum. As a result, the field-and-time 2D profiles were windowed to only include contributions from $|B| > 10$ G (see Fig. S2).

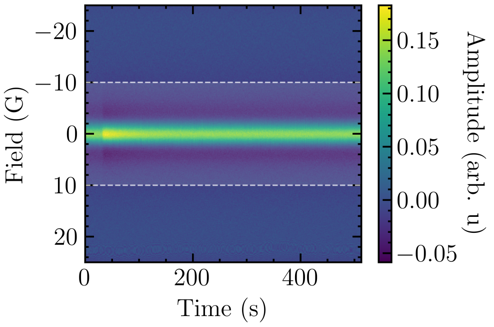


**Fig. S2** Remainder of WT *AsLOV2* data matrix after subtracting a smoothed version of the data. Smoothing $\sigma$ was calculated with an $L$-curve to find the maximum curvature between noise and $\sigma$ value outside of the marked field regions ($|B| > 10$ G) so that the relatively high frequency signal near the peak did not inflate the calculated noise value; $\sigma = 1.9$ G as shown in Fig. S1.

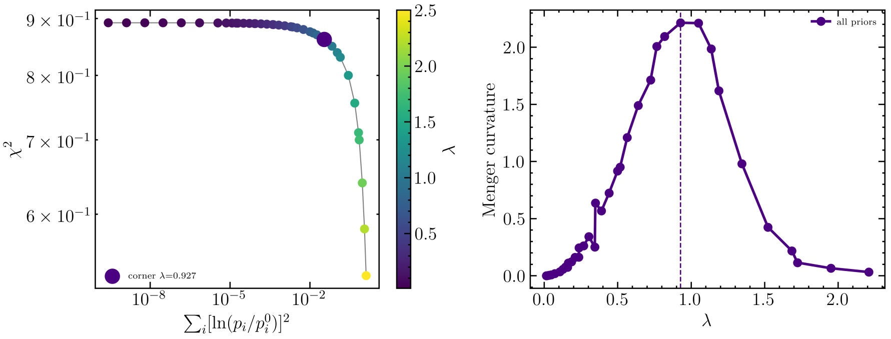


**Fig. S3** (Left) $\chi^2$ and $\lambda$ prior $L$-curve showing the optimal $\lambda$ value to balance the data and priors informing the fit a roughly equal amount. (Right) Menger curvature highlighting the point of maximum curvature between the two regimes; found to be $\lambda = 0.927$ for the WT dataset.

Field and temporal axis window functions were used to reduce the contribution of regions of low signal to the fit. These regions were the wings of the field profiles and the long-time region of the temporal profile, as shown in Fig. S4 and S5.

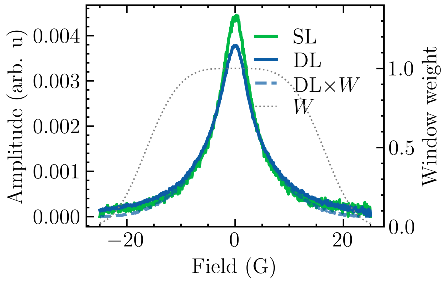


**Fig. S4** Field domain window showing effect of windowing of DL-broadened WT time slice.

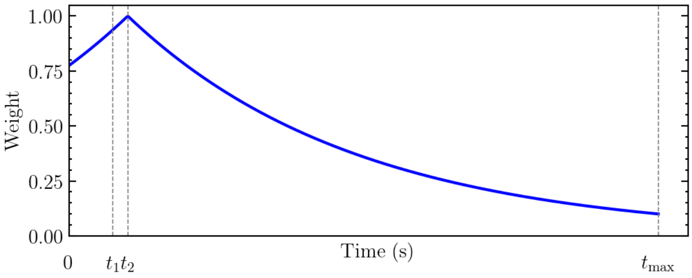


**Fig. S5** Example time domain window depicting coefficient applied to specific time points.

#### S3 Estimation of confidence intervals

Because the residuals are divided by $\sqrt{N_{\text{total}}}$, the $\chi^2$ computed by the fitting code is the standard $\chi^2$ rescaled by the total number of data points, i.e. $\chi^2_{\text{code}} = \chi^2_{\text{standard}}/N_{\text{total}}$. The standard single-degree-of-freedom threshold $\Delta\chi^2_{\text{standard}} \leq 3.84$ [19, 20] therefore becomes

$$\Delta\chi^2_{\text{code}} \leq \frac{3.84}{N_{\text{total}}}. \tag{S1}$$

However, the field-domain data are highly correlated, so there are far fewer than $N_{\text{total}}$ statistically independent measurements. Replacing $N_{\text{total}}$ with $N_{\text{eff}}$, the effective number of independent data points, gives the corrected threshold [**?** ]

$$\Delta\chi^2_{\text{code}} \leq \frac{3.84}{N_{\text{eff}}}. \tag{S2}$$

In practice the threshold is written as

$$\Delta\chi^2_{\text{code}} \leq \frac{3.84\,\chi^2_{\text{min}}}{N_{\text{eff}}} \tag{S3}$$

rather than Eq. (S2) directly. For properly normalized data and a single degree of freedom, a good fit returns $\chi^2_{\text{min}} \approx 1$; here the values were larger than 1 due to imperfections in the model, specifically baseline distortion on the high field side, and the center of the fitted components not reaching the full peak amplitude of the raw data. The $\chi^2_{\text{min}}$ factor corrects for systematic deviation (see Figs. S4 and S5) or otherwise varied $\chi^2$ across datasets. What is left, then, is to estimate $N_{\text{eff}}$. For an autocorrelated dataset,

$$N_{\text{eff}} \approx \frac{N}{1 + 2\sum_{k=1}^{\infty} \rho_k}, \tag{S4}$$

where the denominator is the autocorrelation length [**?** ]. Applying this calculation to the raw data would give erroneously long distances due to the single (highly correlated) underlying EPR spectrum. To mitigate this and estimate the number of "independent samples" in a single time slice, the data was smoothed along the field axis using a Gaussian filter of width $\sigma$ and the smoothed data was subtracted from the raw data, leaving an approximation of the high-frequency (dominant) noise. The center of the EPR spectra remained structured after smoothing, and was therefore windowed out of the noise calculation (only values with $|B| > 10\,\text{G}$ were used, see Fig. S2). The window is shown in Fig. S2. The autocorrelation of the *noise*, then, was calculated for each slice along the time axis and their mean was used to generate the system-wide autocorrelation at each lag, $k$. The optimal $\sigma$ that preserved noise while removing signal was calculated using an $L$-curve (see Fig S1). Rather than sum to infinity, which would reduce the length after the autocorrelation dips below zero, the sum only kept terms up to the first zero crossing (see Fig. S6). This analysis was only applied to the field axis. For the temporal axis, a similar treatment resulted in a roughly white-noise autocorrelation length of approximately one sample, greatly inflating $N_{\text{eff}}$ and thereby strongly shrinking the CIs to unrealistic bounds ($< 0.1\%$ range around LSQ optimum). To conservatively (over-)estimate the confidence intervals, an $N_{\text{eff}}^{\text{time}}$ value was determined from the model itself, which implied that there were only three kinetic regimes: pre-illumination, during illumination (unfolding), and post-illumination (refolding). Therefore,

$$N_{\text{eff}} = N_{\text{eff}}^{\text{time}} * N_{\text{eff}}^{\text{field}} \tag{S5}$$

$$\approx 3 * \frac{N}{1 + 2\sum_{b=1}^{\infty} \rho_b}, \tag{S6}$$

where index $b$ denotes a sum over the magnetic field axis. Using these values, the 95% CI bounds were calculated by starting at the LSQ minima for a given parameter, walking outward in both directions from that minima, and re-fitting using the LSQ routine with all other parameters. This was repeated over a range of possible points for each parameter; if the threshold $\Delta\chi^2_{\text{code}} > \frac{3.84\chi^2_{\text{min}}}{N_{\text{eff}}}$ was crossed, the precise crossing was calculated with linear interpolation between the two points above and below the threshold. Running this algorithm in both directions defined the bounds of the CI. If a sufficient bound was not found for one direction, it was reported as a † in the main text.

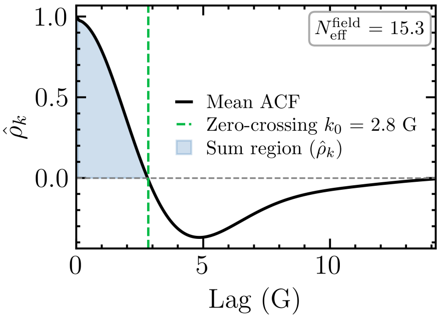


**Fig. S6** Autocorrelation at lag $k$ for WT *AsLOV2* dataset at ∼20 °C used to calculate $N_{\text{eff}}^{\text{field}}$.

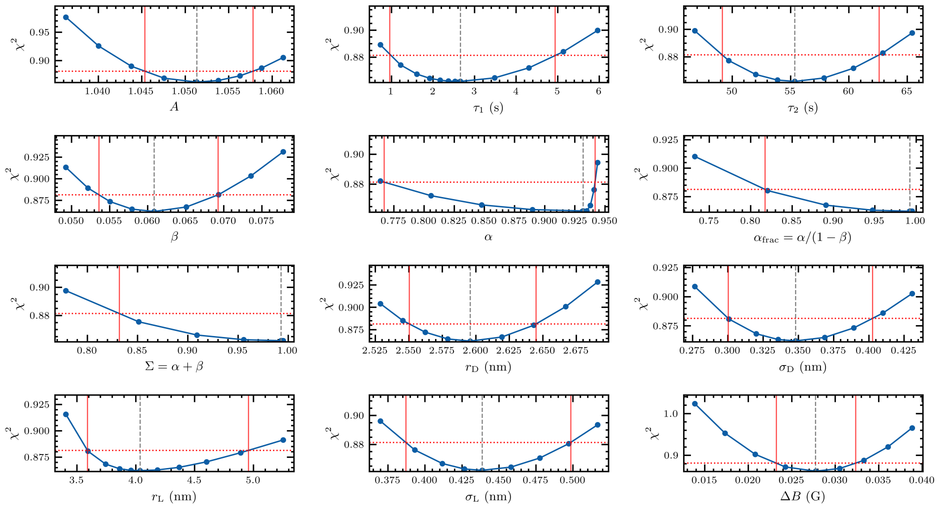


**Fig. S7** $\chi^2$ profiles used to calculate 95% confidence intervals for each of the ten free parameters in the fit. Each parameter was held fixed at varying values (across each $x$-axis) and the fit was re-optimized to compare fit quality as each parameter varied independently. $\chi^2$ thresholds are shown with horizontal dashed red lines and computed 95% CI bounds are shown with vertical dashed red lines.

The bands for Fig. **??** were generated by randomly sampling the (assumed independent) distributions generated by the width of each fit parameter's confidence intervals. This was repeated 10,000 times and the bands represent the range of the 2.5-to-97.5 percentile outcomes. This treatment is an approximation, as the parameters correlated, which is not accounted for by the independent sampling. However, it provides an approximate visual esimate for the 95% confidence intervals for the fit. A representative skew-asymmetric Gaussian distribution is shown in Fig. S9.

For each fit parameter, the confidence-interval sampling distribution was taken to be a continuous, mode-normalized skew-asymmetric ("two-piece") Gaussian,

$$A(z;\mu,\sigma,r) = \frac{2}{\sqrt{2\pi}\,\sigma\,(r+1)} \begin{cases} \exp\left(-\dfrac{(z-\mu)^2}{2\sigma^2}\right), & z > \mu, \\ \exp\left(-\dfrac{(z-\mu)^2}{2r^2\sigma^2}\right), & z \le \mu, \end{cases} \tag{S7}$$

where $\mu$ is the LSQ-optimal parameter value, $\sigma$ is the standard deviation above $\mu$, and $r$ rescales the standard deviation below $\mu$; $A$ is continuous and normalized [21].

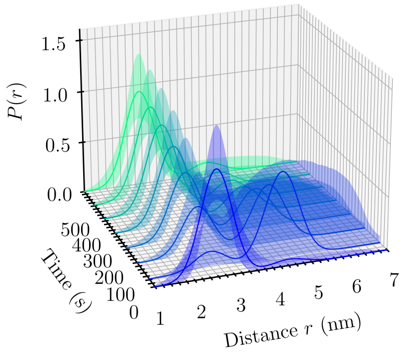


**Fig. S8** Summed Gaussian distance distributions as returned by the best fit of the model to the TiGGER data of N414Q *AsLOV2* at 20 °C. Eight of roughly 500 frames are shown. The sample was illuminated between frames one and two. Light-activated unfolding and refolding is shown, resulting in a large shift in relative near/far populations. Confidence intervals were generated by Monte Carlo sampling skew-asymmetric Gaussian distributions with the 95% CI as 1.96$\sigma$ (see S.I. Fig. S9).

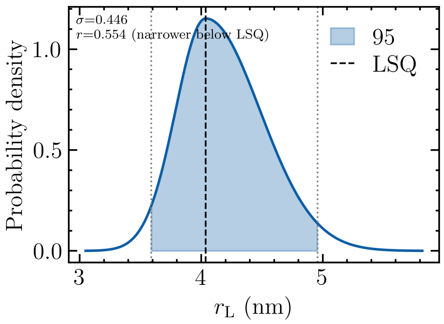


**Fig. S9** Skew-asymmetric profile for WT $r_L$ [21]. $r$ was calculated so that the 1.96$\sigma$ values of this profile were equal to the 95% CIs as shown in Fig. S7.

[2] D.A. Pearlman, D.A. Case, J.W. Caldwell, W.S. Ross, T.E. Cheatham, S. DeBolt, D. Ferguson, G. Seibel, P. Kollman, AMBER, a package of computer programs for applying molecular mechanics, normal mode analysis, molecular dynamics and free energy calculations to simulate the structural and energetic properties

of molecules. Computer Physics Communications **91**(1-3), 1–41 (1995). https://doi.org/10.1016/0010-4655(95)00041-D. URL https://linkinghub.elsevier.com/retrieve/pii/001046559500041D
